# Role of Proinsulin Self-Association in Mutant *INS* gene-induced Diabetes of Youth

**DOI:** 10.1101/786970

**Authors:** Jinhong Sun, Yi Xiong, Xin Li, Leena Haataja, Wei Chen, Saiful A. Mir, Rachel Madley, Dennis Larkin, Arfah Anjum, Balamurugan Dhayalan, Nischay Rege, Nalinda D. Wickramasinghe, Michael A. Weiss, Pamela Itkin-Ansari, Randal J. Kaufman, David A. Ostrov, Peter Arvan, Ming Liu

## Abstract

Abnormal interactions between misfolded mutant and wild-type (WT) proinsulin in the endoplasmic reticulum (ER) drive the molecular pathogenesis of Mutant-*INS*-gene induced Diabetes of Youth (MIDY). How these abnormal interactions are initiated remains unknown. Normally, proinsulin-WT dimerizes in the ER. Here, we suggest that the normal proinsulin-proinsulin contact surface, involving the B-chain, contributes to dominant-negative effects of misfolded MIDY mutants. Specifically, we find that proinsulin Tyr-B16, which is a key residue in normal proinsulin dimerization, helps confer dominant-negative behavior of MIDY mutant proinsulin-C(A7)Y. Substitutions of Tyr-B16 with ether Ala, Asp, or Pro in proinsulin-C(A7)Y each decrease the abnormal interactions between the MIDY mutant and proinsulin-WT, rescuing proinsulin-WT export, limiting ER stress, and increasing insulin production in β-cells and human islets. This study reveals the first evidence indicating that noncovalent proinsulin-proinsulin contact initiates dominant-negative behavior of misfolded proinsulin, pointing to a novel therapeutic target to enhance bystander proinsulin export for increased insulin production.

## Introduction

Proinsulin misfolding, endoplasmic reticulum (ER) stress, and decreased β-cell mass are key features of β-cell failure and diabetes in the autosomal-dominant diabetes known as Mutant-*INS*-gene induced Diabetes of Youth (MIDY) (1–3). All MIDY patients carry a mutation in one of two *INS* alleles (3–6), although, in principle, only one normal *INS* allele should be sufficient to maintain normoglycemia (7). The underlying mechanism of MIDY is thought to involve abnormal interactions of misfolded mutant proinsulin with bystander proinsulin-WT in the ER, impairing the ER export of proinsulin-WT and decreasing production of bioactive mature insulin (8–10). However, to date, how these interactions are initiated remains largely unknown. Interestingly, the clinical spectrum of MIDY ranges from neonatal-onset severe insulin deficient diabetes to relatively late-onset mild diabetes (1, 2), suggesting that different proinsulin mutants may behave differently in their folding processes, as well as their ability to interact and obstruct bystander proinsulin-WT in the ER.

In this study, we report that a proline (Pro) substitution for tyrosine residue at the 16^th^ position of the proinsulin B chain (Tyr-B16) [i.e., proinsulin-Y(B16)P] results in dramatic proinsulin misfolding in the ER, and yet is unable to interact with bystander proinsulin-WT in the ER, and is unable to impair the ER export of proinsulin-WT or to block its ability to form mature insulin. Interestingly, Y(B16)P disrupts the main α-helix in the proinsulin B-domain. Normally, proinsulin uses this B-domain α-helix in order to homodimerize in the ER (11). Thus, we hypothesize that cross-dimerization between MIDY mutant and proinsulin-WT may be required to initiate their abnormal interaction that impairs the intracellular transport of proinsulin-WT, resulting in insulin deficiency that leads to diabetes (12).

Structural evidence strongly implicates Tyr-B16 as a key residue for normal proinsulin dimerization (13). With this in mind, we have mutagenized this residue to either aspartic acid [Y(B16D)] Asp) or alanine [Y(B16)A]; both substitutions were entirely innocuous when introduced into proinsulin-WT. However, when these substitutions were introduced into a MIDY causing *Akita* proinsulin-C(A7)Y, they both could function as an intra-allelic suppressor, blocking abnormal interactions with proinsulin-WT; alleviating dominant-negative behavior on proinsulin-WT intracellular trafficking; and increasing insulin production in pancreatic β-cell lines and human islets. This study provides insight that protecting the proinsulin-proinsulin contact surface might be a potential druggable target to limit abnormal proinsulin interactions and prevent/delay the development and progression of MIDY.

## Results

In the ER, proinsulin dimerization (11, 14) involves nonpolar association of apposed B-chain residues 8-29 (encompassing the central B9-B19 α-helix), which continue to remain associated in mature insulin (15, 16). The insulin crystal structure highlights that B16Y contributes more surface area to the dimerization interface than any other residue (Fig. 1A, B), and a substitution of this residue yields a high potency monomeric insulin (13). We therefore engineered both two-chain and single-chain insulin derivatives bearing the Y(B16)D substitution, both of which showed impaired dimerization in dilute solution as judged by size exclusion chromatography and 1D NMR spectroscopy, respectively (Fig. S1A, B). To determine the role of TyrB16 in proinsulin folding and dimerization in the ER, we expressed proinsulin-Y(B16)D and Y(B16)A in 293T cells, and demonstrated these substitutions affect neither proinsulin ER oxidative folding (Fig. 1C-D) nor ER export (Fig. 1E-F), suggesting that impaired dimerization alone does prevent the intracellular trafficking of proinsulin. However, Y(B16)P that disrupted the proinsulin B9-B19 α-helix indeed impaired formation of proper disulfide bonds and blocked secretion of proinsulin (Fig. 1C-F), suggesting that the central α-helix of the proinsulin B-chain indeed plays an important role in proinsulin folding in the ER.

**Fig. 1.**
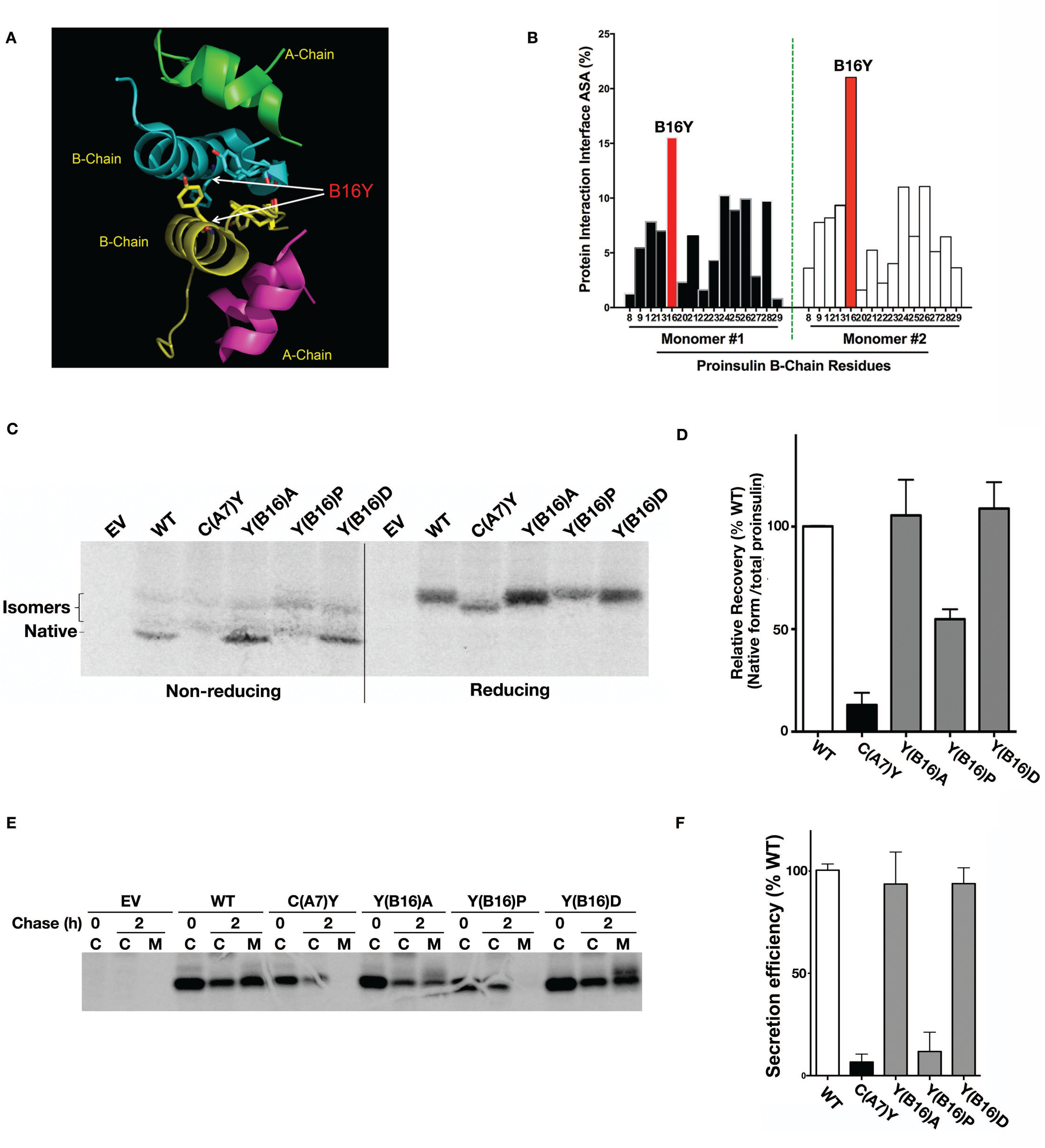
Proinsulin B-chain tyrosine-16 plays an important role in proinsulin-proinsulin contact, folding and secretion. **A.** Ribbon and stick diagram from the insulin crystal structure showing the aromatic side chains of B-chain Tyr-16 near the center of the B-chain—B-chain contact surface. **B.** Residues contributing to the accessible surface area (ASA) at the site of contact between two insulin molecules (B-chain residues B9-B29), identified from the insulin crystal structure using the Protein-Protein Interaction Server. **C and D.** 293T cells transfected with plasmids encoding proinsulin-WT (“WT”), C(A7)Y, Y(B16)A, Y(B16)D, or Y(B16)P were labeled with ^35^S-amino acids for 30 min without chase. The newly synthesized proinsulin was immunoprecipitated with anti-insulin followed by analysis by tris-tricine-urea-SDS-PAGE under both nonreducing and reducing conditions. Native proinsulin (correct disulfide pairing) recovered under nonreducing conditions and total proinsulin recovered under reducing conditions, from at least three independent experiments, were quantified and shown in Fig. 1D. The relative recovery of native proinsulin in the cells expressing PI-WT was set to 100%. **E and F**. 293T cells transfected with plasmids encoding proinsulin-WT (“WT”), C(A7)Y, Y(B16)A, Y(B16)D, or Y(B16)P were labeled with ^35^S-amino acids for 30 min followed by 0 or 2 h chase, as indicated. The cells (“C”) were lysed and the chase media (“M”) collected. All samples were analyzed by immunoprecipitation with anti-insulin, SDS-PAGE under reducing conditions and phosphorimaging (panel E), quantified in panel F.

We hypothesize that the proinsulin-proinsulin contact surface initiates the attack by misfolded MIDY proinsulin mutants on co-expressed proinsulin-WT. We therefore decided to explore the role of proinsulin-Y(B16) in the interactions between the MIDY-causing “*Akita*” proinsulin-C(A7)Y and proinsulin-WT. In 293 cells we co-expressed untagged human proinsulin-WT with Myc-tagged human proinsulin-WT or C(A7)Y [Myc-PI-C(A7)Y] with or without Y(B16)D substitution (Fig. 2A, lanes 4-6 and 4’-6’). By nonreducing SDS-PAGE and anti-proinsulin immunoblotting, the overexpression of either untagged proinsulin-WT (PI-WT) or Myc-tagged proinsulin-WT (Myc-PI-WT) resulted in detectable formation of homotypic disulfide-linked complexes (Fig. 2A, lanes 2 and 3). Recent evidence indicates that *Akita* proinsulin is similarly predisposed to form a ladder of aberrant disulfide-linked complexes even at low expression levels (17–19), and *Akita* proinsulin can heterotypically co-recruit proinsulin-WT into these aberrant complexes (10).

**Fig. 2.**
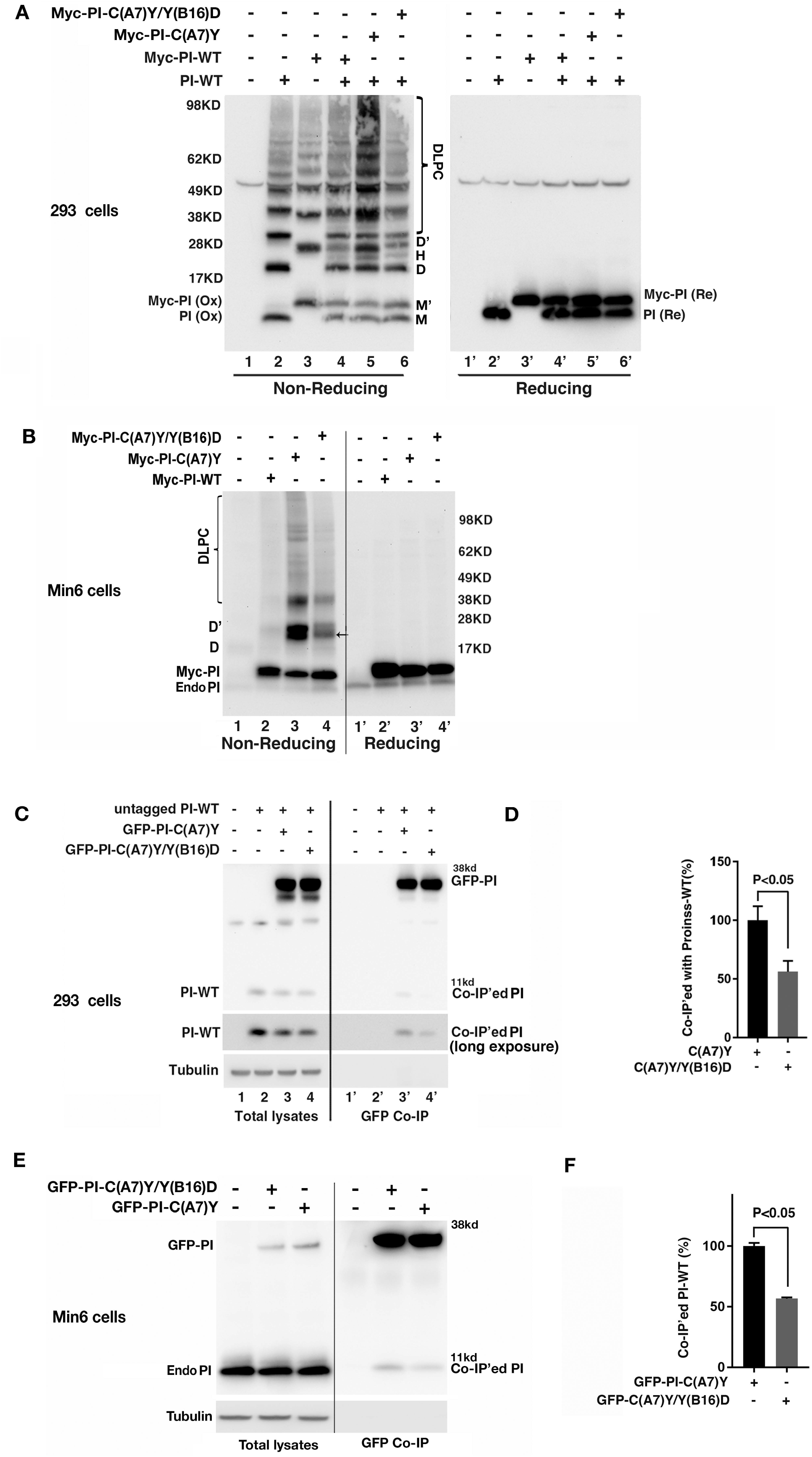
Substitution of Tyr to Asp at proinsulin B16 limits abnormal interactions between wild-type proinsulin and diabetes causing C(A7)Y mutant proinsulin. **A**. 293T cells were transfected with plasmids encoding untagged proinsulin-WT (PI-WT), Myc-tagged proinsulin WT (Myc-PI-WT), or proinsulin mutants as indicated. Oxidative folding, disulfide-linked proinsulin dimers (D refers to homo-dimers formed by un-tagged PI, and D’ refers to homo-dimers formed by Myc-PI, and H refers to cross-dimers formed by untagged PI with Myc-PI), and higher-molecular weight disulfide-linked proinsulin complexes (DLPC) were analyzed under nonreducing (oxidized, Ox) conditions. The total amount of PI-WT and Myc-PI-WT or mutants were analyzed under reducing (Re) condition. **B**. 293T cells were co-transfected with plasmids encoding untagged proinsulin-WT (“PI-WT”), or GFP-tagged proinsulin bearing the C(A7)Y mutation (“GFP-PI-C(A7)Y”), or GFP-tagged proinsulin bearing both the C(A7)Y mutation as well as an intragenic suppressor mutation, Y(B16)D (“GFP-PI-C(A7)Y/Y(B16)D”) at a DNA ratio of 1:1. At 48 h post-transfection, the cells were lysed in a co-IP buffer and immunoprecipitated with anti-GFP. Immunoprecipitated GFP-PI and co-precipitated PI-WT were resolved by 4-12% NuPage under reducing conditions, transferred to the nitrocellulose membrane, and blotted with anti-human proinsulin that can recognize both un-tagged human PI and GFP-PI. **C and D**. Min6 cells transfected with plasmids encoding GFP-PI-C(A7)Y or GFP-PI-C(A7)Y/Y(B16)D were lysed in a co-IP buffer and immunoprecipitated with anti-GFP followed. The endogenous proinsulin (PI) in the total lysate, and that co-IP’ed with anti-GFP, were resolved in 4-12% NuPage under reducing condition, transferred to the nitrocellulose membrane, and blotted with anti-mouse proinsulin, which can recognize both endogenous mouse PI and GFP-PI.

Upon co-expression, although monomeric forms of Myc-PI-WT, C(A7)Y, and C(A7)Y/Y(B16)D were comparable, Myc-PI-C(A7)Y clearly formed more disulfide-linked dimers (untagged PI dimers and Myc-tagged dimers were marked as D and D’, respectively), heterodimers (marked as H), and higher molecular weight proinsulin complexes (Fig 2A, compare lanes 4 and 5). Upon treatment with the reducing agent dithiothreitol (DTT), all of these complexes collapsed into a single band of reduced untagged or Myc-tagged-PI, indicating that the bands seen upon nonreducing SDS-PAGE are disulfide-mediated proinsulin dimers and cross-dimers, as well as disulfide-linked proinsulin complexes (“DLPC”). Importantly, the formation of these abnormal proinsulin complexes was notably decreased in cells expressing the C(A7)Y/Y(B16)D double mutant (Fig. 2A, compare lanes 5 and 6).

To examine whether these disulfide-mediated proinsulin interactions occur in β-cells, we expressed Myc-PI-WT or mutants in Min6 β-cells. As shown in Fig. 2B, Myc-PI-C(A7)Y not only formed significantly more abnormal disulfide-linked proinsulin dimers (marked as D’) compared with that of Myc-PI-WT, but also formed disulfide-linked cross-dimers with endogenous proinsulin (noted by an arrow). The high molecular weight DLPC were also increased in the cells expressing Myc-PI-C(A7)Y (Fig. 2B, compare lanes 2 and 3). However, once again, introducing the intra-allelic Y(B16)D significantly decreased these abnormal proinsulin complexes (Fig. 2B, compare lanes 3 and 4). Together, these results suggest that limiting proinsulin-proinsulin interactions by introducing Y(B16)D can decrease formation of abnormal disulfide-linked proinsulin complexes between proinsulin-C(A7)Y and proinsulin-WT.

To further confirm this point, we expressed GFP-tagged proinsulin-C(A7)Y with or without Y(B16)D substitution in both 293 cells and Min6 cells. We found that Y(B16)D inhibits co-precipitation of proinsulin-WT by GFP-PI-C(A7)Y in both 293 cells (Fig. 2C-D) and Min6 cells (Fig. 2E-F) under steady-state conditions (Fig. 2C-F) and for newly synthesized proinsulin molecules (Fig. S2). Thus, the Y(B16)D substitution decreases the strength of interaction of MIDY proinsulin with proinsulin-WT in the ER.

Next, we asked whether the decreased interaction of MIDY proinsulin-C(A7)Y with proinsulin-WT could also alleviate the dominant-negative effect on proinsulin-WT secretion. We first examined folding and secretion of proinsulin-C(A7)Y with or without B16 substitutions, and found that introduction of Asp (D), Ala (A), or Pro (P) had no effect on the impaired disulfide maturation of proinsulin-C(A7)Y and did not rescue secretion of the MIDY mutant (which was retained intracellularly; Fig. 3A-B). We then checked whether B16 substitutions could limit the effect of proinsulin-C(A7)Y on bystander proinsulin-WT. We co-expressed mouse proinsulin-C(A7)Y ± Y(B16)D or Y(B16)P substitutions with human proinsulin-WT in HEK293T cells. We used a species-specific immunoassay to follow secretion of the co-expressed human proinsulin-WT in the presence of mutant proinsulin. Indeed, the expression of mouse proinsulin-C(A7)Y blocked the secretion of proinsulin-WT by ∼70% (Fig. 3C), consistent with previous reports (9, 10, 20, 21). However, the Y(B16)D substitution [or Y(B16)P] functioned as an intra-allelic suppressor of the dominant-negative effect of proinsulin-C(A7)Y on the secretion of co-expressed proinsulin-WT (Fig. 3C).

**Fig. 3.**
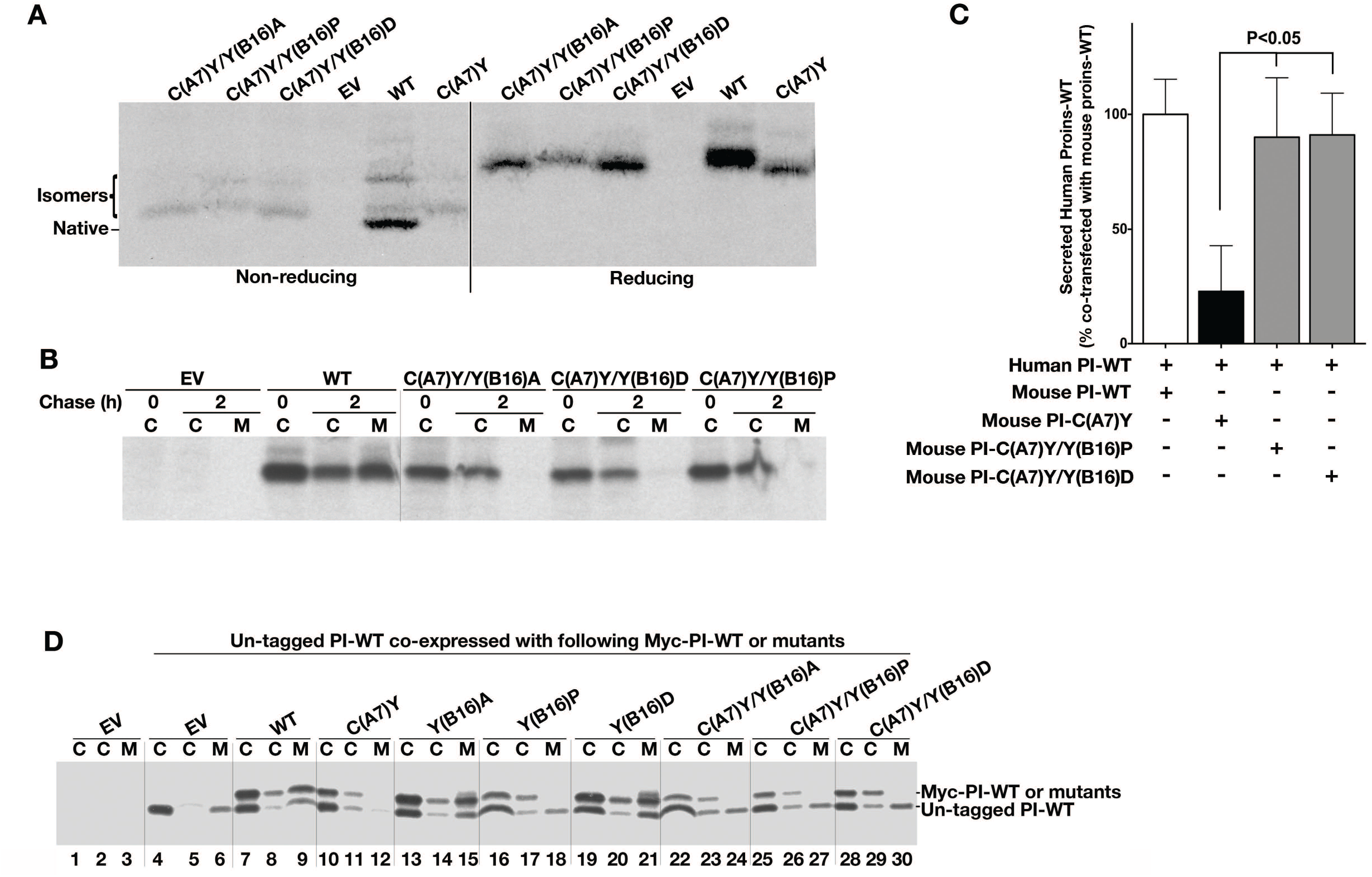
Mutations at proinsulin B16 do not improve the folding of *Akita* proinsulin-C(A7)Y but alleviate its dominant-negative effect on co-expressed proinsulin-WT. **A.** 293T cells co-transfected with the indicated plasmids were labeled with ^35^Met/Cys for 30 min without chase. Biosynthesis and folding of newly synthesized PI-WT and mutants were analyzed by tris-tricine-urea-SDS-PAGE under both nonreducing and reducing conditions. **B**. 293T cells transfected and pulse-labeled as in A were chased for 0 or 2 h. The cells (“C”) were lysed and chase media (“M”) collected and analyzed by immunoprecipitation with anti-insulin followed by SDS-PAGE under reducing conditions, followed by phosphorimaging. **C.** 293T cells were co-transfected with a plasmid encoding human proinsulin-WT (human PI-WT) and either mouse proinsulin-WT (mouse PI-WT) or mutants as indicated. The secretion of human PI-WT in the presence of mouse PI-WT or mutants was measured using human-PI-specific RIA (mean ± SD from at least 3 independent experiments). **D**. 293T cells were co-transfected as Fig. 3C. At 48 h post-transfection, the cells were labeled with ^35^Met/Cys for 30 min with 0 or 3 h chase. The cell lysates and chase media were immunoprecipitated with anti-insulin. The secretion efficiency of un-tagged PI-WT in the presence or absence of Myc-PI-WT or mutants were analyzed under reducing conditions, followed by phosphorimaging.

These immunoassay results were then confirmed using pulse-chase experiments. Once again, proinsulin-Y(B16)A and proinsulin-Y(B16)D were secreted normally, in parallel with the secretion of co-expressed proinsulin-WT (Fig. 3D, lanes 15 or 21). Further, both misfolded mutants proinsulin-C(A7)Y and proinsulin-Y(B16)P were fully blocked in their secretion; however, the Akita proinsulin-C(A7)Y conferred a clear dominant-negative effect on co-expressed proinsulin-WT that was not apparent for proinsulin-Y(B16)P (Fig. 3D, lanes 12 *vs* 18), indicating that not all misfolded proinsulin mutants have the ability to block secretion of bystander proinsulin-WT. Moreover, we found that neither proinsulin-C(A7)Y/Y(B16)A nor C(A7)Y/Y(B16)D could block the secretion of co-expressed proinsulin-WT (Fig. 3D, lanes 12 *vs* 24 or 30), indicating that these Tyr-B16 substitutions reversed the dominant-negative effect of *Akita* mutant proinsulin.

To further investigate effect of B16 substitutions on endogenous proinsulin in β-cells, we transfected GFP-tagged proinsulin-WT or mutants, and used three independent approaches to examine trans-dominant effects. First, using confocal immunofluorescence, we found that while GFP-proinsulin-WT and Y(B16)D showed a punctate insulin secretory granule pattern, proinsulin-C(A7)Y appeared in a typical ER pattern and caused a profound loss of endogenous insulin (Fig. 4A white arrows). However, whereas the double mutant proinsulin-C(A7)Y/Y(B16)D also appeared to be retained in the ER, it did not detectably decrease endogenous insulin in β-cells (Fig. 4A bottom, white arrowheads). Second, using human insulin specific radioimmunoassay, we confirmed that introducing Y(B16)D in proinsulin-C(A7)Y alleviated the trans-dominant effect of the MIDY mutant as demonstrated by increased insulin production (Fig. 4B). Further, in human islets, adenoviral expression of mouse proinsulin-C(A7)Y significantly decreased human proinsulin and insulin content, whereas comparable expression of the C(A7)Y/Y(B16)D mutant (Fig. S3) again demonstrated intra-allelic suppression of the dominant-negative effect of the MIDY mutant on both human proinsulin and insulin (Fig. 4D). Finally, the enhanced ER export of proinsulin-WT correlated with decreased ER stress in β-cells co-expressing proinsulin-C(A7)Y versus C(A7)Y/Y(B16)D, as measured in a transcriptional BiP promoter reporter assay (Fig. 4D). Taken together, these results strongly suggest that Tyr(B16), which is a major component of the proinsulin-proinsulin interaction surface, is involved in the physical interaction between MIDY mutant proinsulin and proinsulin-WT, and this interaction is necessary for the mutant protein to block proinsulin-WT exit from the ER, impair insulin production, trigger ER stress, and promote the development of diabetes.

**Fig. 4.**
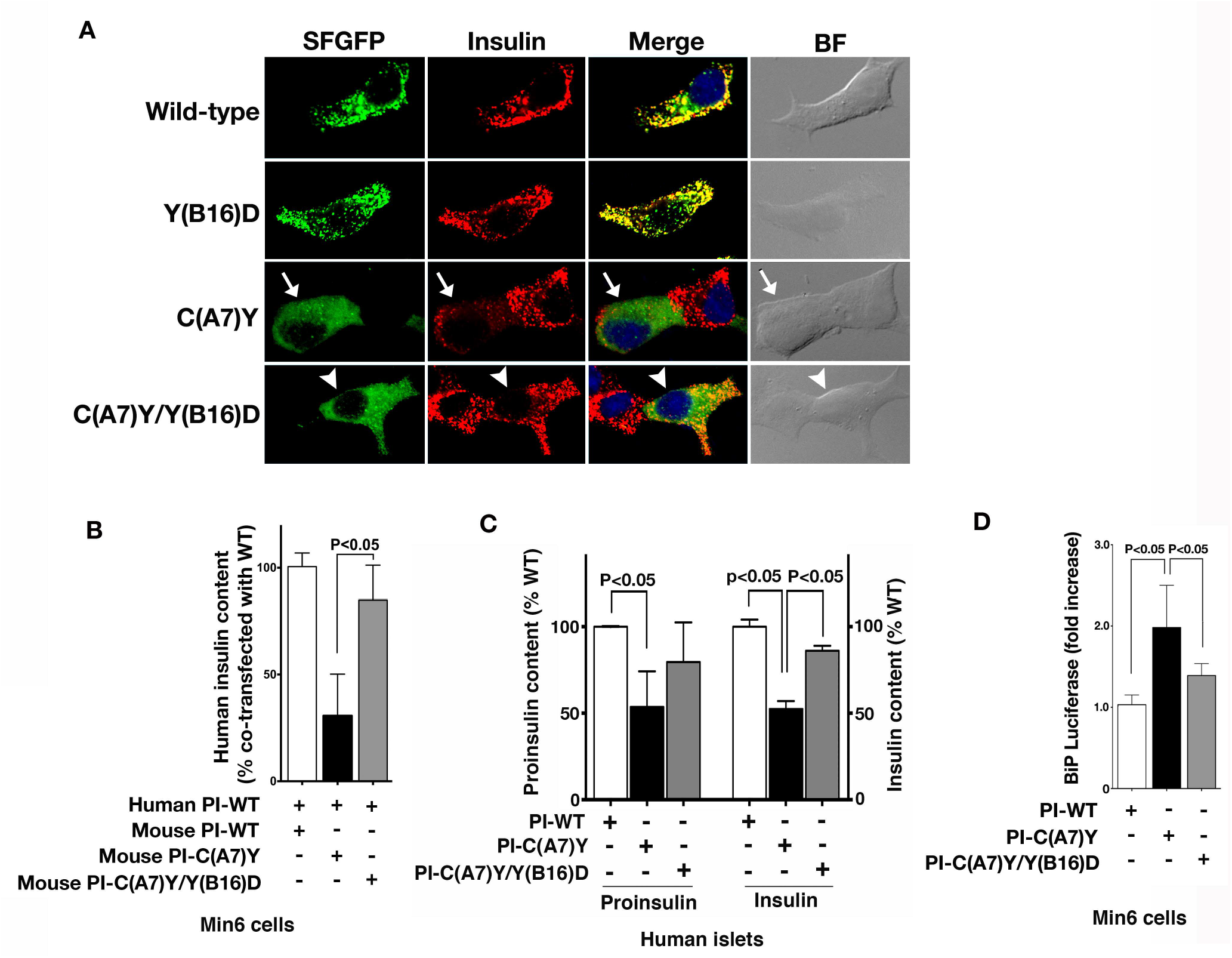
Proinsulin B16 substitutions function as an intragenic suppressor of the dominant-negative effects of MIDY proinsulin-C(A7)Y, and alleviate β-cell ER stress. **A.** INS1 cells were transfected with plasmids encoding GFP-tagged proinsulin-WT and mutants as indicated. At 48 h post-transfection, the cells were fixed and permeabilized. **B.** Confocal immunofluorescence microscopy was performed after double-labeling with anti-GFP (green) and anti-insulin (red). Whereas the cells expressing GFP-C(A7)Y exhibited a diminution of endogenous insulin staining (middle panel, arrows), in the cells expressing a GFP-C(A7)Y/Y(B16)D endogenous insulin was restored (bottom panel, arrowheads). **B**. MIN6 cells were co-transfected with plasmids encoding human proinsulin-WT with either mouse proinsulin-WT or mutants at a DNA ratio of 1:3. At 48 h post-transfection, human insulin content in the transfected cells was measured using human insulin specific RIA (mean ± SD from at least 3 independent experiments). **C.** Human islets were transduced with adenoviruses expressing Myc-tagged mouse proinsulin-WT, C(A7)Y, or C(A7)Y/Y(B16)D, as indicated. At 48 h post infection, human insulin content in the infected human islets was measured by human insulin specific ELISA. Human proinsulin content was measured by densitometry of Western blotting (Fig. S3). The relative contents of human proinsulin and insulin in human islets infected with mouse proinsulin-WT were set to 100%. **D.** MIN6 cells were triple-transfected with plasmids encoding firefly luciferase driven by a BiP-promoter; CMV-renilla luciferase; and either proinsulin-WT or mutants at a DNA ratio of 1:2:4. At 48 h post-transfection, luciferase activities in transfected cells were measured as described in Methods (mean ± SD from at least 3 independent experiments).

## Discussion

To date, more than 50 insulin gene mutations have been reported to cause MIDY, an autosomal dominant form of diabetes (also known as MODY10) (8, 12). More than 60% of the identified mutant alleles have been predicted or experimentally confirmed to impair oxidative folding of proinsulin in the ER (1, 22). Importantly, these misfolded MIDY mutants not only fail to exit from the ER, but also physically attack co-expressed proinsulin-WT, impairing the folding and ER export of co-expressed proinsulin-WT, decreasing insulin production, and initiating insulin-deficient diabetes (12, 20, 23). Heretofore, the molecular mechanism underlying these dominant-negative effects have been completely unknown. As the proinsulin-proinsulin contact surface might possibly engage even before completion of the folding of individual proinsulin monomers, the present work focuses attention on this surface as a possible site initiating the attack of misfolded mutant proinsulin onto bystander proinsulin in the ER.

In this study, we have found at least two substitutions of the key Tyr-B16 residue that have no observable impact on proinsulin monomer folding or ER export, but can limit proinsulin-proinsulin contact and thereby can decrease abnormal interactions and alleviate trans-dominant effects of observed in MIDY. It is interesting to note that among the roughly 30 known proinsulin sites/residues that trigger MIDY, none has yet been described that introduces a charged or highly polar residue into the B9-B19 α-helix. We postulate that such substitutions may be relatively ineffective in propagating misfolding onto bystander proinsulin-WT molecules. This hypothesis is supported by data from the proinsulin-Y(B16)P mutant, which disrupts the B9-B19 α-helix to cause severe proinsulin misfolding and ER retention (Fig. 1C-FF) but cannot impose a dominant-negative effect on bystander proinsulin-WT (Fig. 3D). These data support the notion that proinsulin-proinsulin contact is an early folding event and initiates the trans-dominant effects of MIDY mutants.

Increasing evidence indicates that proinsulin misfolding and ER stress not only are the molecular basis of MIDY, but may also play an important role in the development and progression of type 1 and type 2 diabetes (2, 24–27). Increased proinsulin misfolding has been recently reported in β-cells with either defects in the ER folding environment (28–30), or defective ER export machinery (18), or proinsulin oversynthesis due to insulin resistance in rodent and human type 2 diabetes islets (19). These data suggest that proinsulin misfolding is an early event in the progression to type 2 diabetes. As the most abundant protein synthesized in β-cells, proinsulin is predisposed to misfold (12, 20, 31) from which it may propagate this misfolding onto bystander proinsulin (8–10). This becomes an important driving force for β-cell ER stress and insulin deficiency (29, 32–34). The data in this report provide the first evidence pointing to the idea that initial proinsulin-proinsulin B-chain contact may be a site worth targeting, to block cross-dimerization of misfolded proinsulin with proinsulin bystanders, enhance insulin production, and offer a new approach to preventing β-cell insulin deficiency and diabetes (Fig. 5).

**Fig. 5.**
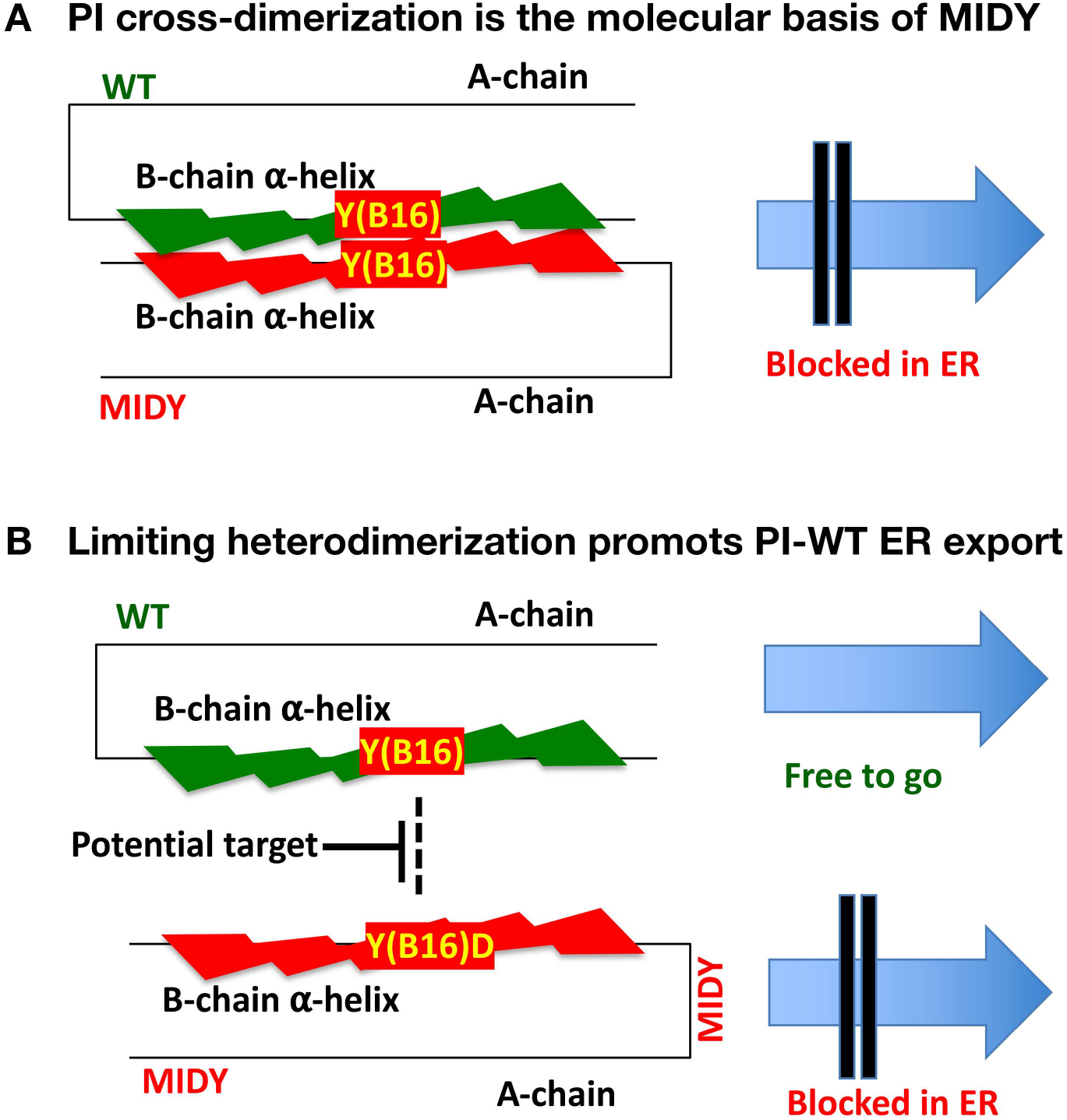
A working model of misfolded mutant proinsulin attacking the dimerization surface of bystander proinsulin-WT as a control point for proinsulin ER export and insulin production. **A.** Proinsulin (PI) forms dimers in the ER. In the β-cells that co-express proinsulin-WT (green) and misfolded MIDY mutant proinsulin (red), the proinsulin dimerization surface of the mutant can abnormally interact with that of the co-expressed bystander proinsulin-WT, which impairs the folding and ER export of proinsulin-WT. Proinsulin TyrB16 plays a key role in proinsulin dimerization in the ER. **B.** Protecting the proinsulin dimerization surface may sever as a potential theropeutic target to limit *trans*-dominant effects of misfolded mutant proinsulin, alleviating ER stress and β-cell failure.

## Acknowledgements

This work was supported by NIH R01-DK48280, R01-DK88856, R24-DK110973, and National Natural Science Foundation of China 81830025 and 81620108004. We acknowledge support from the Michigan Diabetes Research Center Morphology Core (NIH P30 DK020572), and support of the Protein Folding Diseases Initiative of the University of Michigan.

---

**Fig. S1.**
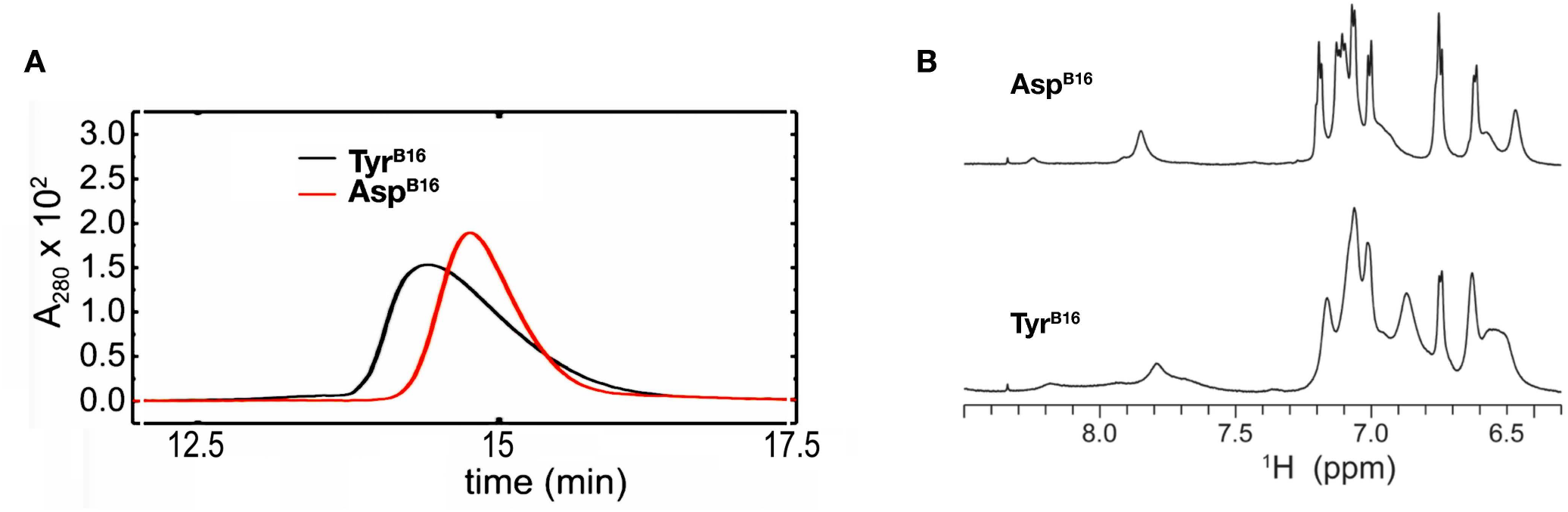
Substitution of Tyr to Asp in the insulin B-chain 16^th^ residue inhibits dimerization. **A.** Two-chain insulin analogues bearing either Tyr^B16^ (black line) or Asp^B16^ (red line) were synthesized (see Methods), and retention times from 0.6 mM solutions were measured in a zinc-free buffer by size-exclusion chromatography (SEC). Quantification of these data demonstrate that the Asp^B16^ analogue forms at least three times more monomer than the Tyr^B16^ analogue. **B.** Single-chain Tyr^B16^-DesDi insulin analogue or Asp^B16^-DesDi insulin analogue were synthesized (49 residues, see Methods). The aromatic region from 1D ^1^H NMR spectra at 700 MHz and 25° C reveal resonance broadening due to self-association of Tyr^B16^-DesDi single-chain insulin analog (below), which is notably mitigated in the Asp^B16^-DesDi single-chain insulin analog (above).

**Fig. S2.**
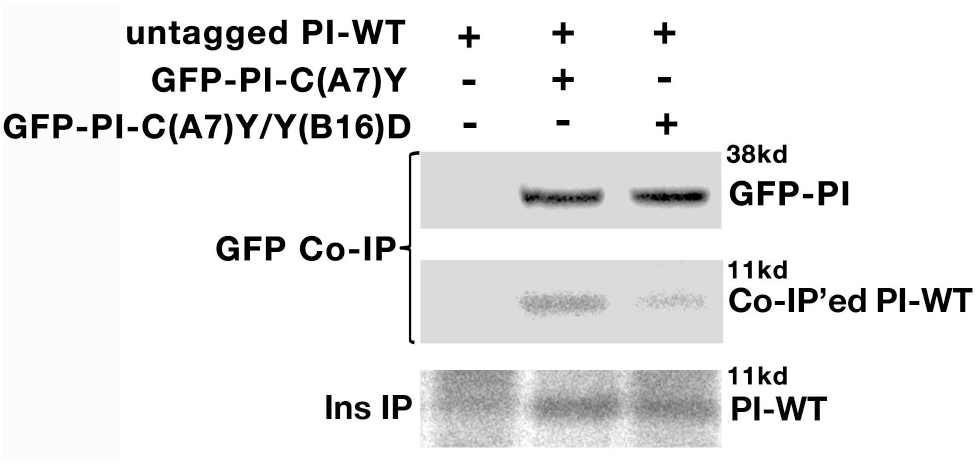
A Y(B16)D mutation decreases abnormal interactions between newly synthesized proinsulin-WT with co-expressed proinsulin-C(A7)Y. 293T cells were co-transfected with plasmids encoding untagged proinsulin-WT and GFP-PI-C(A7)Y or GEP-PI-C(A7)Y/Y(B16)D. The cells were labeled with ^35^Met/Cys for 30 min without chase. The cells were lysed and split into two halves, followed by IP with anti-insulin (bottom panel) and anti-GFP (upper and middle panels), respectively. The co-IP’ed untagged PI-WT is shown in the middle panel. The data are representative of three replicate experiments.

**Fig. S3.**
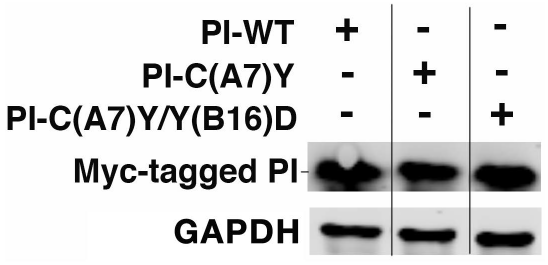
Adenoviral-expression of Myc-tagged proinsulin-WT, C(A7)Y, and C(A7)Y/Y(B16)D in human islets. Human islets infected with adenovirus expressing either Myc-tagged mouse proinsulin-WT, or C(A7)Y, or C(A7)Y/Y(B16)D. This figure (a control for Figure 4C) establishes equivalent viral infection in each sample, as shown by Western blot for the Myc-tagged murine proinsulins, and equivalent protein levels in the human islet cell lysate samples, normalized both by protein assay and protein loading control (GAPDH).

